# Gene expression noise randomizes the adaptive response to DNA alkylation damage in E. coli

**DOI:** 10.1101/551911

**Authors:** Stephan Uphoff

## Abstract

DNA damage caused by alkylating chemicals induces an adaptive response in *Escherichia coli* cells that increases their tolerance to further damage. Signalling of the response occurs through methylation of the Ada protein which acts as a damage sensor and induces its own gene expression through a positive feedback loop. However, random fluctuations in the abundance of Ada jeopardize the reliability of the induction signal. I developed a quantitative model to test how gene expression noise and feedback amplification affect the fidelity of the adaptive response. A remarkably simple model accurately reproduced experimental observations from single-cell measurements of gene expression dynamics in a microfluidic device. Stochastic simulations showed that delays in the adaptive response are a direct consequence of the very low number of Ada molecules present to signal DNA damage. For cells that have zero copies of Ada, response activation becomes a memoryless process that is dictated by an exponential waiting time distribution between basal Ada expression events. Experiments also confirmed the model prediction that the strength of the adaptive response drops with increasing growth rate of cells.

## INTRODUCTION

The accurate detection and repair of DNA damage is crucial for genome stability and cell survival. In addition to constitutively expressed repair pathways, cells employ DNA damage responses that activate DNA repair factors in the presence of DNA damage. The fidelity of the DNA repair system relies on a series of processes: sensing the presence of DNA damage or DNA damaging agents, inducing a DNA damage response, and correctly repairing lesions. Cell strains with genetic defects that impair the function of any of these processes show sensitivity to DNA damage, elevated mutation rates, and genome instability. However, even in fully repair-proficient cell strains the accuracy of the DNA repair system is fundamentally limited by the stochastic nature of the molecular interactions involved (1, 2): For example, proteins that signal or repair DNA damage perform a random target search and therefore have a finite chance of overlooking lesions (3–7). The repair process itself can also be error-prone and cause mutations, loss, or rearrangements of genetic material (8–12). Traditionally, research has focused on genetic defects and such “intrinsic errors” in DNA repair – i.e. errors that are inherent to the repair mechanism and thus occur with the same probability in all cells of a population.

By comparison, less attention has been given to “extrinsic variation” in the DNA repair system – i.e. fluctuations in protein abundances that may affect the repair capacity of individual cells. Gene expression noise is ubiquitous (13, 14) and difficult for cells to suppress (15). Feedback gene regulation can establish bimodal distributions so that subpopulations of cells maintain distinct states of gene expression for long times. Whereas many biological processes are robust to a certain level of noise, even transient variation in the capacity of a cell to repair DNA damage can have severe and potentially irreversible consequences (16–18). For instance, cells that transiently express too little of a damage sensor protein may be unable to signal DNA damage efficiently, leading to mutations or cell death. But there may also be evolutionary benefits to heterogeneity and occasional errors in the DNA repair system when cells are facing selective pressure (19–21).

The adaptive response to DNA alkylation damage in *Escherichia coli* is a case where gene expression noise appears to cause significant cell-to-cell heterogeneity in DNA repair capacity (18, 17). Alkylated DNA lesions block DNA replication and transcription and can lead to mutations (22, 10). The adaptive response is regulated by the Ada protein, a DNA methyltransferase that directly repairs methylated phosphotriester and O^6^MeG lesions by transferring the methyl group from the DNA onto its own cysteine residues (23–25). These reactions are irreversible and turn the methylated Ada protein (meAda) into a transcriptional activator of the genes *ada*, *alkB*, *alkA*, and *aidB* that are involved in the repair or prevention of DNA alkylation damage (26). The response causes a positive feedback amplification of Ada expression that renders cells more tolerant to further damage. Surprisingly, the timing of response activation varies drastically across genetically identical cells even at saturating levels of DNA damage (18). The very low abundance of Ada before DNA damage treatment appears to be responsible for this variation. In particular, single-molecule imaging showed that stochastic Ada expression results in a subpopulation of cells that does not contain a single Ada protein and therefore cannot sense the presence of DNA alkylation damage. Without induction of the adaptive response, the insufficient repair capacity of these cells increases mutation rates to the same level as in mutant cells in which the *ada* gene has been deleted (17).

These surprising observations call for a quantitative model to pinpoint the noise source responsible for heterogeneity in the adaptive response. Quantitative models have been key to our current understanding of gene expression noise and its important functions in diverse biological processes (13, 27–30), including DNA damage signalling and repair (16, 31–34). Here, I capitalized on time-lapse microscopy data for the adaptive response that were recorded for a large range of damage conditions in hundreds of single *E. coli* cells (18). The direct measurement of key observables and parameters allowed construction of a quantitative model of the core Ada regulation. The proposed model is remarkably simple, yet accurately reproduces experimental observations – both the cell average as well as the stochastic behaviour of single cells. The model also predicts cell responses after different experimental perturbations. No additional post-hoc noise term was required in our model but propagation of basic Poisson fluctuations alone was sufficient to explain the observed cell-to-cell variation in response activation. These results establish that intrinsic noise in the basal expression of the *ada* gene is solely responsible for the stochastic nature of the adaptive response. The model also predicts that the strength of the response should be inversely related to the growth rate of cells, which was confirmed in experiments.

## MATERIALS AND METHODS

### Experimental data

The construction of the model was based on experimental data described in reference (18). Briefly, the adaptive response was monitored in live *E. coli* AB1157 cells carrying a functional fusion of Ada to the fast-maturing fluorescent protein mYPet (35) that is expressed from the endogenous chromosomal locus, thus maintaining native expression levels. Single cells growing continuously inside the “mother machine” microfluidic device (36) were treated with the DNA methylating agent methyl methanesulfonate (MMS) and Ada-mYPet fluorescence was measured using time-lapse microscopy in multiple fields of view at 3-minute intervals. Fluorescence intensities were calculated from the average pixel intensities within the segmented cell areas. To correct for the background fluorescence, the intensity before MMS treatment was subtracted on a per-cell basis.

Additional experiments (data in Fig. 5) used the same microfluidic imaging setup and acquisition parameters as described in our previous work (17). The only difference was that cell growth rates were varied using minimal medium supplemented either with glucose or glycerol as carbon sources.

### Ada response model

The structure of the model is based on previous genetic and biochemical characterization of the adaptive response (23–25). Key to the model is a positive feedback loop in which DNA damage-induced methylation of Ada creates meAda, which acts as a transcriptional activator for the *ada* gene. The chemical kinetics of the model can be described as a system of ODEs according to the diagram in Figure 1A:

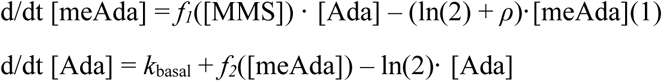

**Fig. 1.**
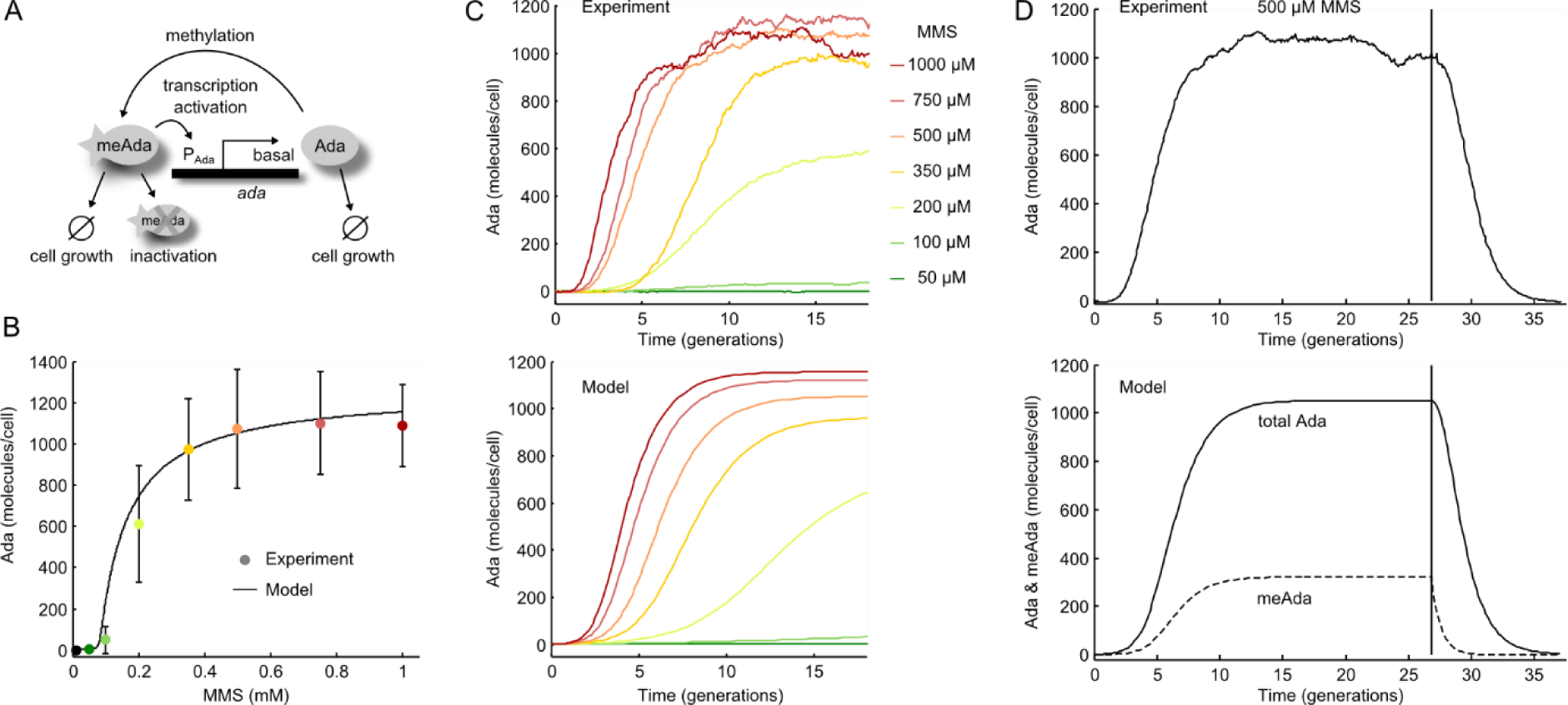
Deterministic model of the adaptive response. **(A)** Schematic of the model: Ada is expressed at a low basal level in undamaged cells. MMS treatment creates DNA methylation damage that converts Ada to meAda. ada transcription is activated by meAda binding to the P_Ada_ promoter. Both Ada and meAda molecules are diluted due to cell growth and division. Additionally, meAda gets inactivated by degradation or auto-repression. **(B)** Average steady-state expression of Ada after constant treatment with different doses of MMS for 20 cell generations (mean ± standard deviation). The curve shows the analytical steady-state solution of the total Ada abundance. **(C)** Average response induction dynamics when MMS was added at time 0 and Ada abundance was measured by time-lapse fluorescence microscopy. The model curves were generated by numerically solving the rate equations for different MMS concentrations. **(D)** Average response induction and deactivation dynamics upon addition and removal of 500 µM MMS. The vertical line indicates the time of MMS removal. For the model, total Ada (Ada + meAda + inAda) and meAda levels are shown as solid and dashed lines, respectively.

Self-methylation of Ada in the presence of DNA methylation damage generates meAda molecules with a rate proportional to the MMS concentration: *f*_1_([MMS]) = *k*_me_ · [MMS].

In the absence of DNA methylation damage, the *ada* gene is expressed at a constant basal rate *k*_basal_ from the P_Ada_ promoter. Transcription of the *ada* gene is induced to rate *k*_ind_ when meAda binds to the P_Ada_ promoter with an association rate *k*_on_ and dissociation rate *k_off_* in a non-cooperative manner (37):

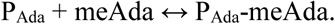

In the deterministic model, Ada is produced according to the fraction of time that the P_Ada_ promoter is bound by meAda:

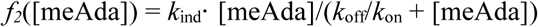

where *k*_ind_ is the fully induced production rate at saturating amounts of meAda.

Production of Ada and meAda molecules is counteracted by dilution due to exponential cell growth. When time is expressed in units of generation times, the dilution rate is equal to ln(2). In addition to dilution, our model also includes loss of meAda at a constant rate *ρ*. This feature was required to match the rapid deactivation of Ada expression upon MMS removal that we observed in experiments and as previously suggested (38, 39). The equation governing the concentration of the inactivated Ada species is:

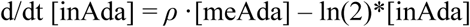

This formulation of the model approximates protein expression as one reaction where transcription and translation are described with a single production rate constant. This reduces the number of free parameters of the model and allows direct comparison of the experimental observables (i.e. Ada-mYPet proteins) with the variables in the model. The simplification is valid when protein expression follows first-order kinetics with a single rate-limiting step. This is consistent with the complete lack of *ada* expression bursting in our experiments (18), and a short half-life and low translation efficiency of *ada* mRNAs (40, 41).

### Steady-state solution

Setting equations (1–3) to zero gives the abundances of Ada and meAda at steady-state. These can be expressed as the solution of a quadratic equation:

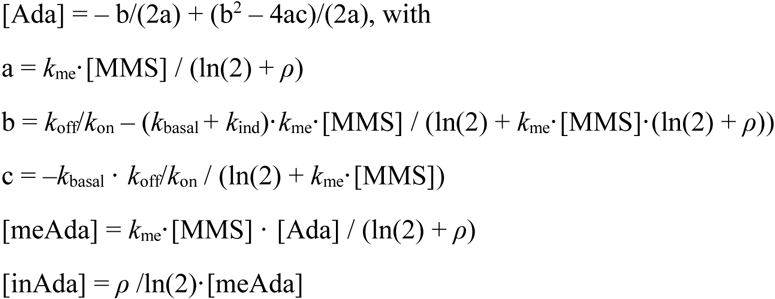

The total Ada level corresponding to the measured Ada-mYPet fluorescence is given by the sum [Ada] + [meAda] + [inAda].

### Numerical solution

The time-dependent solution of the model equations was numerically obtained using the ode45 solver in MATLAB.

### Stochastic simulation

We simulated time-traces of Ada expression in single cells using a custom implementation of Gillespie’s algorithm in MATLAB (42). To this end, the equations (1-3) of the deterministic model were expressed as elementary unimolecular or bimolecular reactions. Gillespie’s algorithm assumes memoryless kinetics, which is appropriate for transitions between discrete chemical states where the system is defined entirely by its present state (Markov process). Stochasticity arises due to the discreteness of the states of the system (i.e. the integer number of molecules in the cell) with spontaneous random transitions given by the elementary reactions of the system. At a given time point, the waiting time until the next transition is drawn from an exponential distribution with an expectation value given by the inverse of the sum of all rates exiting that state (i.e. the rates of molecule production, conversion, and loss). Which of the possible transitions occurs is then chosen randomly with probabilities according to the relative rates of the reactions. Initial molecule numbers were drawn from a Poisson distribution defined by the basal expression rate (see Fig. 2A).

**Fig. 2.**
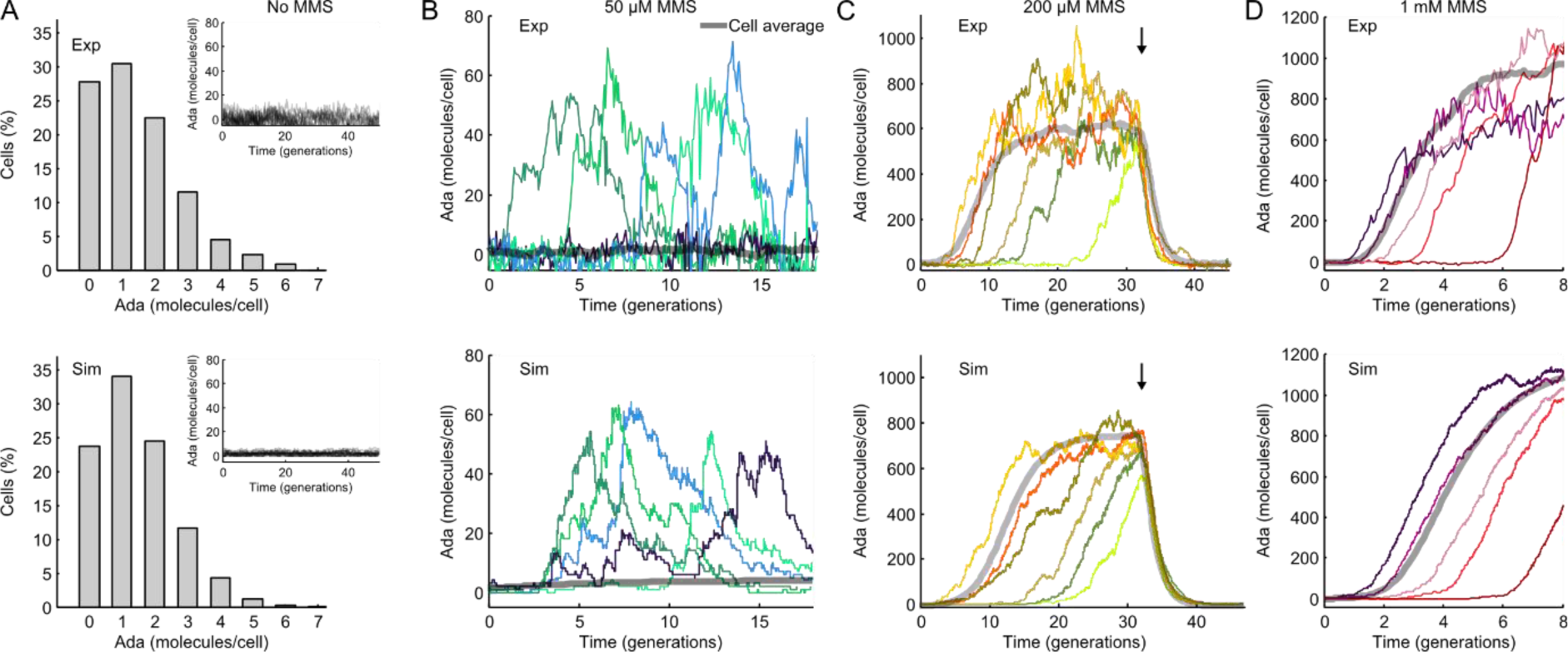
Stochastic dynamics of the adaptive response in single cells. **(A)** Distribution of the number of Ada molecules per cell without MMS treatment. The experimental data (top) is from single-molecule counting experiments (18). Simulated data (bottom) were drawn from a Poisson distribution with a mean of 1 per cell. Inset: Single-cell time-traces of Ada expression without MMS treatment. **(B)** Example time-traces showing random unsynchronized pulses of Ada expression with 50 µM MMS treatment for experimental (top) and simulated data (bottom). The cell-average data is shown in grey. **(C-D)** Stochastic activation of the Ada response with 200 µM and 1 mM MMS treatment. Example time-traces for experimental (top) and simulated data (bottom) are shown. The arrow indicates the time when MMS was removed. The cell-average data is shown in grey.

### Model parameters

Parameters were either obtained by direct experimental measurement (18), or by matching the model output to experimental observations (Table 1). One set of parameters was used for all the deterministic or stochastic model realizations in this paper. The cell generation time of 42 min in supplemented M9 glucose medium at 37°C (or 75 min in M9 glycerol, Fig 5) was obtained directly by timing cell division events in the microfluidic experiments. To directly compare Ada abundances between experiment and model, fluorescence intensity units were converted to molecule concentrations as described previously (18, 43). To account for incomplete fluorescent protein maturation and presence of photobleached mYPet molecules, we estimated that the detection efficiency of fluorescent Ada-mYPet is 80% of the total abundance.

**Table 1.**
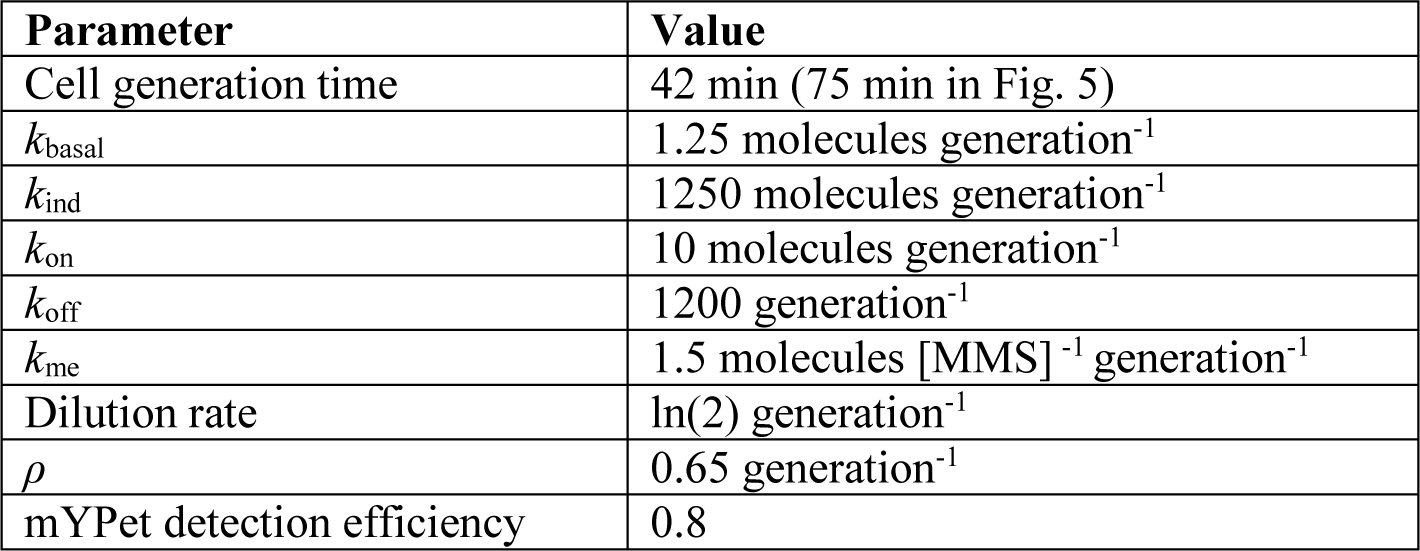
Parameter values used for all plots in this paper.

| Parameter | Value |
| --- | --- |
| Cell generation time | 42 min (75 min in Fig. 5) |
| $k_{\text{basal}}$ | 1.25 molecules generation<sup>-1</sup> |
| $k_{\text{ind}}$ | 1250 molecules generation<sup>-1</sup> |
| $k_{\text{on}}$ | 10 molecules generation<sup>-1</sup> |
| $k_{\text{off}}$ | 1200 generation<sup>-1</sup> |
| $k_{\text{me}}$ | 1.5 molecules [MMS]<sup>-1</sup> generation<sup>-1</sup> |
| Dilution rate | $\ln(2)$ generation<sup>-1</sup> |
| $\rho$ | 0.65 generation<sup>-1</sup> |
| mYPet detection efficiency | 0.8 |

## RESULTS

### A quantitative model of the adaptive response

I first examined whether the proposed model (Fig. 1A) could reproduce the cell-average steady-state expression of Ada after continuous treatment with MMS for 20 cell generations. Experiments showed that Ada expression was very low in the absence of damage treatment and for concentrations below a threshold of 200 µM MMS. A switch-like dose response occurred for concentrations above 350 µM MMS where Ada expression saturated with a very narrow transition region of intermediate Ada expression levels. The analytical steady-state solution of the model accurately reproduced the dose response curve (Fig. 1B).

Imaging cells inside a microfluidic device allowed following the gene expression dynamics of the adaptive response (18). Ada-mYPet expression was measured using time-lapse imaging for hundreds of cells over tens of generations per experiment. Averaging the measured fluorescence signal over all cells at each time point showed that continuous treatment with high MMS concentrations (>350 µM) caused rapid activation of Ada expression within 2 cell generations and steady-state expression was reached within ∼10 generations (Fig. 1C). For lower MMS concentrations (<350 µM), initial response activation was delayed by more than 5 generations and expression reached steady state only after ∼20 generations of treatment. The numerical solution of the model closely matched the measured dynamics (Fig. 1C), using the same set of parameters as for the steady-state analysis. Furthermore, the model confirms that removal of MMS leads to deactivation of the adaptive response (Fig 1D). The abundance of meAda decayed exponentially immediately after MMS removal, whereas Ada remained induced for several generations until meAda levels had diminished. There were no signs of hysteresis effects after MMS removal, as expected for the non-cooperative promoter binding of meAda (37).

### Poisson fluctuations of Ada expression in the absence of DNA damage

Stochastic effects appear to play a key role in the dynamics of the adaptive response at the single-cell level. Single-molecule counting of Ada-mYPet showed that the average production rate in the absence of DNA methylation damage is as low as 1 Ada molecule per cell generation (18). This is equivalent to a population average of 1.4 Ada molecules per cell, given that the loss rate by cell division is ln(2) per cell generation. The distribution of Ada numbers ranged from 0 to ∼6 molecules per cell. Spontaneous induction of higher Ada expression in the absence of MMS treatment was never observed in experiments (Fig. 2A inset). Ada copy numbers were well described by a simulated Poisson distribution when the mean was fixed by the average expression from experiments (Fig. 2A). The integer numbers of Ada molecules can be viewed as discrete cell states and transitions between these states occur with a constant (memoryless) probability given by the average production and loss rates.

Many genes are expressed in bursts, where multiple mRNAs are produced in a short interval and each transcript is translated repeatedly (44), which broadens the expression distributions (45). The close fit of the Poisson distribution demonstrates a lack of expression bursting for Ada, which can be explained by the low translation efficiency and short half-life of *ada* mRNAs (40, 41). It has also been shown that periodic changes in the gene copy number due to DNA replication result in gene expression variation (46, 47). However, because the *ada* gene is located close to the chromosome terminus region the gene is present at a single gene copy throughout most of the cell cycle and therefore expected to show little expression variation over the cell cycle.

### A stochastic model recapitulates single-cell response dynamics after DNA damage treatment

In contrast to the gradual response induction suggested by the numerical solution of the model (Fig. 1D), single-cell measurements revealed significant heterogeneity in *ada* expression after MMS exposure (18). Continuous treatment with low concentrations of MMS (50 - 100 µM) caused stochastic pulses of Ada expression but did not sustain conversion of Ada to the meAda transcription activator (Fig. 2B). Because the pulses are rare, the cell-average expression is close to the low average value predicted by the deterministic version of the model (grey curves in Fig. 2B-D). Intermediate MMS concentrations (200 – 350 µM) resulted in persistent Ada expression once the response was activated, but activation times were extremely broadly distributed across cells (Fig. 2C). Delays of more than 20 generations were frequently observed, a time in which a single cell can grow into a colony of millions. Even at high MMS concentrations (500 µM – 2 mM), activation times differed by multiple generations between cells (Fig. 2D). Contrary to response activation, removal of MMS caused all cells to switch off the adaptive response uniformly (Fig. 2C).

I tested whether the proposed model could explain aspects of the observed cell-to-cell variation. Importantly, the microfluidic imaging system ensures that cells grow under constant identical conditions such that any heterogeneity can be attributed to stochastic processes intrinsic to the cell. I hypothesized that incorporating the discrete nature of molecule numbers and probabilistic reaction kinetics into the model could account for the stochastic response dynamics. This hypothesis was driven by the fact that Poisson fluctuations are especially pronounced for low numbers of molecules as measured for Ada. Moreover, the positive feedback loop of Ada can amplify any initial fluctuations (48).

Stochastic simulations provide a general approach for generating single-cell trajectories that can be directly compared to experimental data (49). To this end, I expressed the model equations as unimolecular or bimolecular elementary reactions and used Gillespie’s algorithm (42) to create probabilistically exact realizations of the proposed model. I used the same parameter values as for the deterministic model. Remarkably, simulated trajectories closely resembled the complex dynamics of the adaptive response in single cells over the whole range of MMS concentrations used in the experiments (Fig. 2). In particular, simulations reproduced the random Ada expression bursts at low MMS as well as the stochastic activation followed by sustained Ada expression at high MMS concentrations. Simulated cell traces also showed uniform deactivation of Ada expression after MMS removal. Importantly, no additional features or noise terms had to be added to the model to achieve these features.

### Poisson noise in basal Ada expression dictates stochastic response delays

For a quantitative comparison of experiments and model simulations, I evaluated the distribution of delay times between addition of MMS and first activation of the adaptive response in single cells (Fig. 3A). The delay time distributions from stochastic simulations of the model closely resembled those from experiments. However, it is evident that the fluctuations in Ada expression after response activation are larger in experiments than in the simulated trajectories (Fig. 2). The additional variation likely reflects “extrinsic noise” (50) due to fluctuations in factors that influence Ada expression but were not included in the model, such as RNA polymerase and ribosome concentrations and variation in the length of the cell cycle. Nevertheless, the perfect match of the simulated and experimental delay time distributions shows that stochasticity in the initial activation time of the response is not influenced by such external noise sources but can be solely attributed to basic Poisson fluctuations in *ada* gene expression.

**Fig. 3.**
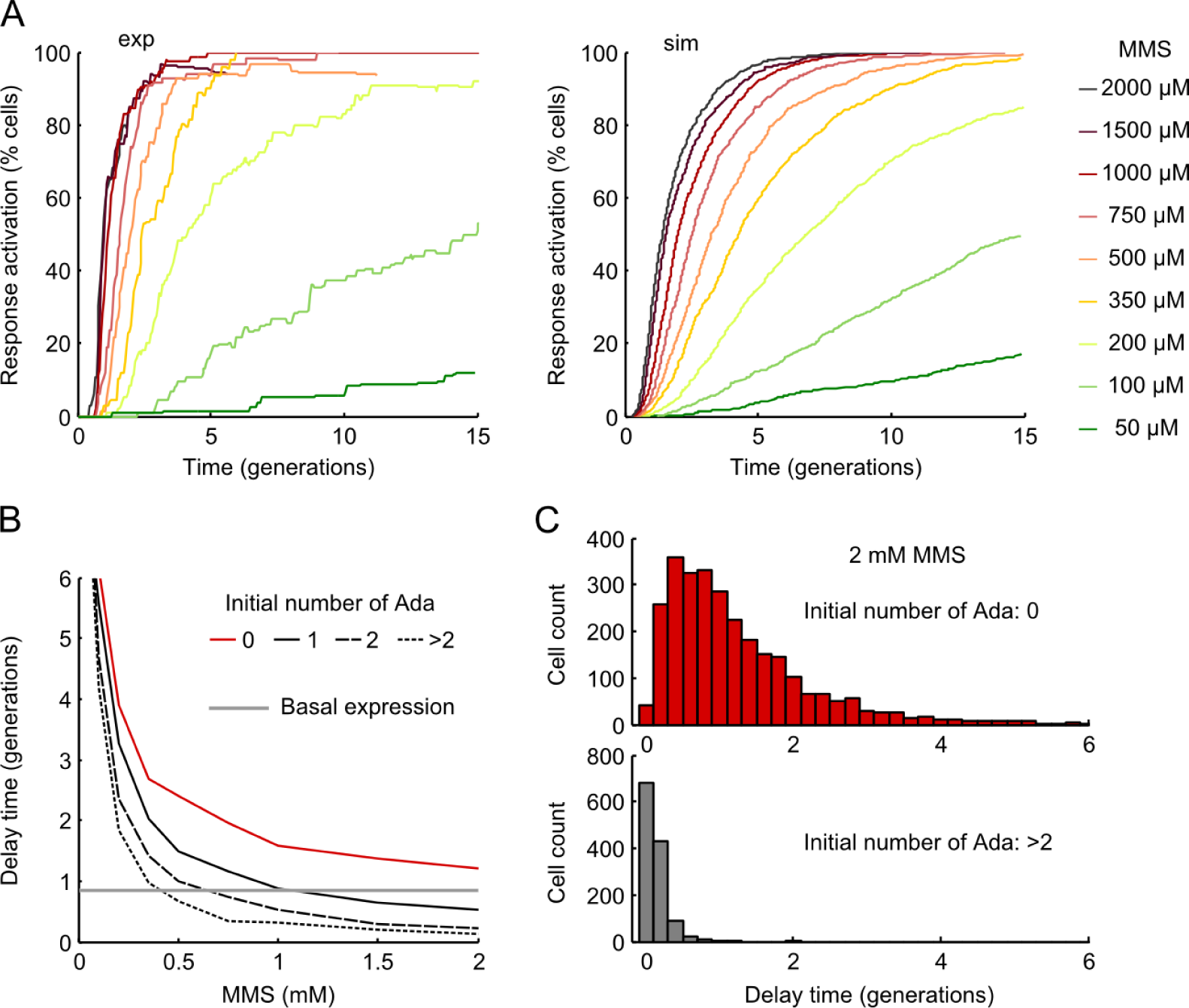
Activation of the adaptive response is delayed by gene expression noise. **(A)** Cumulative distribution showing the percentage of cells that activated the adaptive response after MMS exposure. The activation time corresponds to the time when Ada levels crossed a threshold of 23 molecules/cell (a threshold close to the experimental detection limit). The experimentally measured distributions include between 59 and 149 cells for each MMS concentration (left). Model results are from 1000 independent Gillespie simulations for each MMS concentration (right). Simulated cell trajectories were analysed identical to experimental data. **(B)** Average delay time between MMS addition and generation of the first meAda molecule from 1000 simulated trajectories, conditional on the initial number of Ada molecules at the time of MMS addition. The red curve is for cells that initially had zero Ada molecules; solid black line for 1 Ada molecule, dashed black line for 2 Ada molecules, dotted black line for more than 2 Ada molecules. The average waiting time between basal expression events is shown in grey. **(C)** Simulated distribution of response delay times after 2 mM MMS treatment for cells with initially zero Ada molecules (red), or more than two Ada molecules (grey).

In particular, noise in the low basal expression of Ada is responsible for a subpopulation of 20-30% of cells that do not contain any Ada molecules (18). These cells are thus unable to activate the auto-regulatory adaptive response until they generate at least one Ada molecule. For simulated data, it is possible to calculate response delay times conditional on the initial number of Ada molecules at the time of MMS exposure. This analysis confirmed that the average delay time between MMS addition and generation of the first meAda molecule converges to zero with increasing MMS concentration only for cells that initially contain one or more Ada molecules (Fig. 3B-C). But for cells lacking any Ada molecules, the average delay time approaches a limit defined by the average waiting time between stochastic basal expression events (Fig. 3B-C). In the model, the basal *ada* production is a zero-order reaction with an MMS-independent rate constant. Thus, response activation for cells without Ada molecules follows a memoryless process with an exponential distribution of delay times (Fig. 3C), as seen in experiments (18).

### Testing the predictive power of the model using experimental perturbations

The predictive power of the model was tested by comparison to experiments in which cells were subjected to perturbations that alter the regulation of the adaptive response in a defined manner (18) (Fig. 4). When cell division was inhibited for 45 minutes using the antibiotic cephalexin prior to MMS treatment, Ada molecules accumulate in cells and activation of the adaptive response becomes uniform in the population (18). In agreement with this, prohibiting loss of molecules for 45 minutes was sufficient to generate a uniform response in the simulations (Fig. 4B). The response was also perturbed genetically by supplementing endogenous Ada-mYPet expression with a plasmid that is present at 1-2 copies per cell and expresses *ada* from the P_Ada_ promoter. The slight overexpression of Ada strongly reduced cell-to-cell variation upon MMS treatment and eliminated the population of cells with a delayed response (18). I modelled this perturbation by duplicating the *ada* gene copy number in the simulations (Fig. 4C). This alteration resulted in uniform response activation as seen in experiments (Fig. 4C). However, the simulations generated higher Ada expression levels than measured experimentally, likely because Ada overexpression is toxic in experiments (18).

**Fig. 4.**
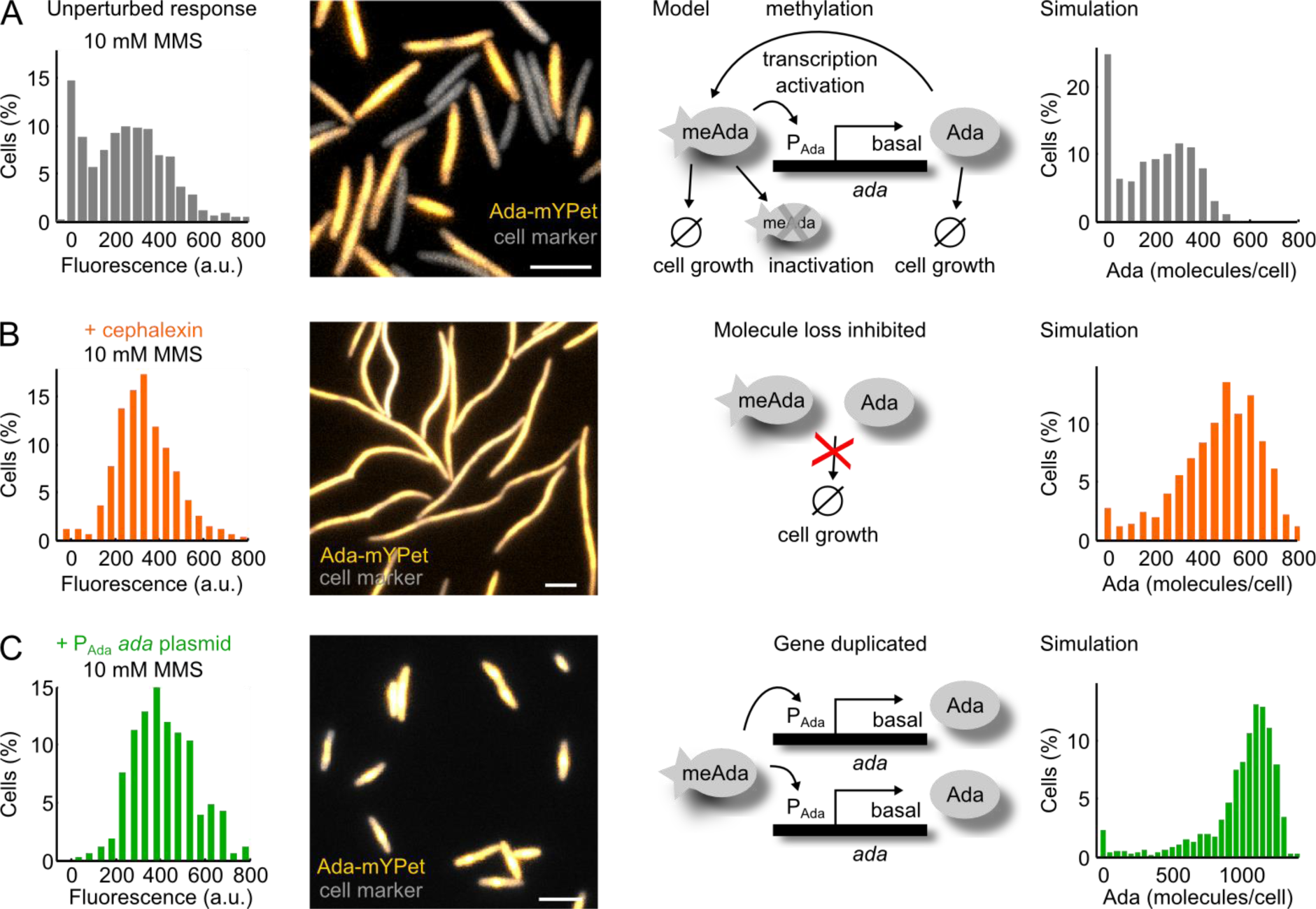
Experimental perturbations to test the model of the adaptive response. The distributions were generated from 1000 simulation repeats. Note that the experiments are shown in fluorescence units (a.u.), while simulation outputs are in molecule numbers. Scale bars: 5 µm. **(A)** Unperturbed Ada-mYPet expression after cell treatment with 10 mM MMS for 1 hour. **(B)** Cephalexin treatment for 45 min before addition of 10 mM MMS for 1 hour. In the simulation, molecule loss was abolished while all other parameters remained unchanged. **(C)** Mild overexpression of Ada by transforming cells that express Ada-mYPet endogenously with a very low copy number plasmid (∼1 per cell) carrying the P_Ada_ promoter and ada gene. Cells were treated with 10 mM MMS for 1 hour before imaging. In the simulation, the ada gene was duplicated while all other parameters remained unchanged.

**Fig. 5.**
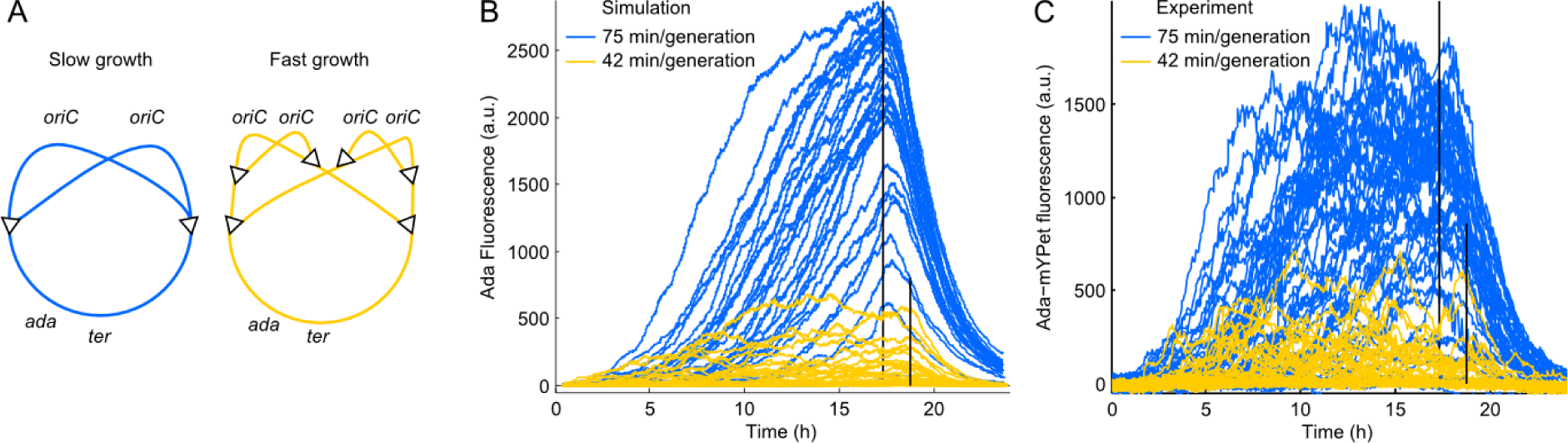
Effect of the cell growth rate on the strength of the adaptive response. **(A)** Schematic showing replication of the circular E. coli chromosome during slow (blue) and fast (yellow) cell growth. Replication initiates at oriC and proceeds bidirectional towards ter (triangles). For fast growth initiation occurs before completion of the previous replication round. The ada gene is located close to ter, so the gene copy number does not increase at faster growth. **(B)** Simulated single-cell traces for slow (blue) and fast (yellow) growth with 100 µM MMS treatment. **(C)** Experimental data with 100 µM MMS treatment for cells growing in minimal medium plus glycerol at 75 min/generation (blue) or in minimal medium plus glucose (yellow) at 42 min/generation. Vertical lines denote times when MMS was removed.

### Effect of cell growth rate on the response strength

The doubling time of *E. coli* in rich growth medium is shorter than the time required to replicate the chromosome. This is achieved by initiating new rounds of replication before completion of the previous round (51). The early duplication of genes close to the replication origin increases their expression proportional to the replication initiation frequency and thus maintains protein abundances at faster growth. Expression of the *ada* gene, however, being located at 49.7 min on the chromosome map in the vicinity of the terminus region, is expected to drop with increasing growth rates (Fig. 5A). This prediction can be tested by the model of the Ada response. I fixed the Ada expression rate while modifying the growth rate, and hence dilution rate. A strong Ada response occurred during slow growth whereas faster growth did not sustain a response at the same MMS concentration (Fig. 5B). I tested this prediction experimentally by growing cells in minimal medium supplemented with glucose or glycerol carbon sources, which lead to 42 min or 75 min generation times, respectively. These measurements confirmed the inverse relation between the growth rate and the strength of the adaptive response (Fig. 5C).

## DISCUSSION

The role of noise in the fidelity of DNA repair has been investigated in eukaryotes, where nucleotide excision repair involves stochastic and reversible assembly of repair factors into large complexes (52, 53). Collective rate control renders the overall repair pathway robust to variation in the abundances of the individual components (34). The situation is opposite for damage signalling by Ada, which acts alone in the regulation of the adaptive response and feedback amplification results in extreme sensitivity to gene expression noise. Remarkably, random variation in the abundance of Ada by just a single molecule was responsible for separating isogenic cells into distinct populations that either induced or failed to induce the DNA damage response. This had important consequences because the lack of a damage response decreased survival and increased mutation in those cells (17). The adaptive response has been described as “a simple regulon with complex features” (24). Instead of attempting to incorporate all mechanistic details, the model described in this article attempts to reduce the *ada* regulation to its central features. For example, methylation of both Cys38 and Cys321 residues in the N- and C-terminal Ada domains is required for optimal activation of the P_Ada_ promoter (54). The model uses only one effective methylation rate and does not distinguish between single or double methylation of Ada. Furthermore, unmethylated Ada has been reported to inhibit meAda-dependent transcription activation (39), a feature that was not explicitly included in the model. The adaptive response also interacts with other cellular responses and processes. For instance, the alternative sigma factor RpoS induces Ada expression upon entry into stationary phase (24), while the SOS response is crucial for initial survival of alkylation damage and contributes to alkylation-induced mutagenesis (17). Considering these simplifications, it is remarkable that the most parsimonious model of the adaptive response not only succeeds in quantitatively reproducing a large spectrum of stochastic single cell dynamics but also in predicting the system’s behaviour after different experimental perturbations.

## ACKNOWLEDGMENTS

I thank Johan Paulsson, David Sherratt, and members of the Uphoff lab for discussions. This work was funded by Sir Henry Wellcome (101636/Z/13/Z) and Sir Henry Dale Fellowships (206159/Z/17/Z), a Wellcome-Beit Prize (206159/Z/17/B), and a Hugh Price Fellowship at Jesus College, Oxford.

The author declares no conflict of interest.

## REFERENCES

1. Uphoff, S., and A.N. Kapanidis. 2014. Studying the organization of DNA repair by single-cell and single-molecule imaging. DNA Repair.

2. Uphoff, S., and D.J. Sherratt. 2017. Single-Molecule Analysis of Bacterial DNA Repair and Mutagenesis. Annu. Rev. Biophys. 46: 411–432.

3. Gorman, J., F. Wang, S. Redding, A.J. Plys, T. Fazio, S. Wind, E.E. Alani, and E.C. Greene. 2012. Single-molecule imaging reveals target-search mechanisms during DNA mismatch repair. Proc. Natl. Acad. Sci. 109: E3074–E3083.

4. Halford, S.E., and J.F. Marko. 2004. How do site-specific DNA-binding proteins find their targets? Nucleic Acids Res. 32: 3040–3052.

5. Kad, N.M., and B. Van Houten. 2012. Dynamics of lesion processing by bacterial nucleotide excision repair proteins. Prog. Mol. Biol. Transl. Sci. 110: 1–24.

6. Stracy, M., M. Jaciuk, S. Uphoff, A.N. Kapanidis, M. Nowotny, D.J. Sherratt, and P. Zawadzki. Single-molecule imaging of UvrA and UvrB recruitment to DNA lesions in living Escherichia coli. Nat. Commun. : in press.

7. Uphoff, S., R. Reyes-Lamothe, F.G. de Leon, D.J. Sherratt, and A.N. Kapanidis. 2013. Single-molecule DNA repair in live bacteria. Proc. Natl. Acad. Sci. U. S. A. 110: 8063–8068.

8. Chen, J., B.F. Miller, and A.V. Furano. 2014. Repair of naturally occurring mismatches can induce mutations in flanking DNA. eLife. 3: e02001.

9. Foti, J.J., B. Devadoss, J.A. Winkler, J.J. Collins, and G.C. Walker. 2012. Oxidation of the Guanine Nucleotide Pool Underlies Cell Death by Bactericidal Antibiotics. Science. 336: 315–319.

10. Fu, D., J.A. Calvo, and L.D. Samson. 2012. Balancing repair and tolerance of DNA damage caused by alkylating agents. Nat. Rev Cancer. 12: 104–120.

11. Giroux, X., W.-L. Su, M.-F. Bredeche, and I. Matic. 2017. Maladaptive DNA repair is the ultimate contributor to the death of trimethoprim-treated cells under aerobic and anaerobic conditions. Proc. Natl. Acad. Sci. : 201706236.

12. Moore, J.M., R. Correa, S.M. Rosenberg, and P.J. Hastings. 2017. Persistent damaged bases in DNA allow mutagenic break repair in Escherichia coli. PLOS Genet. 13: e1006733.

13. Raser, J.M., and E.K. O’Shea. 2005. Noise in Gene Expression: Origins, Consequences, and Control. Science. 309: 2010–2013.

14. Balazsi, G., A. van Oudenaarden, and J.J. Collins. 2011. Cellular decision-making and biological noise: From microbes to mammals. Cell. 144: 910–925.

15. Lestas, I., G. Vinnicombe, and J. Paulsson. 2010. Fundamental limits on the suppression of molecular fluctuations. Nature. 467: 174–178.

16. Paek, A.L., J.C. Liu, A. Loewer, W.C. Forrester, and G. Lahav. 2016. Cell-to-Cell Variation in p53 Dynamics Leads to Fractional Killing. Cell. 165: 631–642.

17. Uphoff, S. 2018. Real-time dynamics of mutagenesis reveal the chronology of DNA repair and damage tolerance responses in single cells. Proc. Natl. Acad. Sci. 115: E6516–E6525.

18. Uphoff, S., N.D. Lord, B. Okumus, L. Potvin-Trottier, D.J. Sherratt, and J. Paulsson. 2016. Stochastic activation of a DNA damage response causes cell-to-cell mutation rate variation. Science. 351: 1094–1097.

19. Bjedov, I., O. Tenaillon, B. Gérard, V. Souza, E. Denamur, M. Radman, F. Taddei, and I. Matic. 2003. Stress-induced mutagenesis in bacteria. Science. 300: 1404–1409.

20. Galhardo, R.S., P.J. Hastings, and S.M. Rosenberg. 2007. Mutation as a Stress Response and the Regulation of Evolvability. Crit. Rev. Biochem. Mol. Biol. 42: 399–435.

21. Yaakov, G., D. Lerner, K. Bentele, J. Steinberger, and N. Barkai. 2017. Coupling phenotypic persistence to DNA damage increases genetic diversity in severe stress. Nat. Ecol. Evol. 1: s41559-016-0016-016.

22. Sedgwick, B. 2004. Repairing DNA-methylation damage. Nat. Rev. Mol. Cell Biol. 5: 148–157.

23. Samson, L., and J. Cairns. 1977. A new pathway for DNA repair in Escherichia coli. Nature. 267: 281–283.

24. Landini, P., and M.R. Volkert. 2000. Regulatory Responses of the Adaptive Response to Alkylation Damage: a Simple Regulon with Complex Regulatory Features. J. Bacteriol. 182: 6543–6549.

25. Sedgwick, B., P.A. Bates, J. Paik, S.C. Jacobs, and T. Lindahl. 2007. Repair of alkylated DNA: Recent advances. DNA Repair. 6: 429–442.

26. Mielecki, D., M. Wrzesiński, and E. Grzesiuk. 2015. Inducible repair of alkylated DNA in microorganisms. Mutat. Res. Rev. Mutat. Res. 763: 294–305.

27. Mettetal, J.T., D. Muzzey, J.M. Pedraza, E.M. Ozbudak, and A. van Oudenaarden. 2006. Predicting stochastic gene expression dynamics in single cells. Proc. Natl. Acad. Sci. 103: 7304–7309.

28. Norman, T.M., N.D. Lord, J. Paulsson, and R. Losick. 2013. Memory and modularity in cell-fate decision making. Nature. 503: 481–486.

29. Patange, O., C. Schwall, M. Jones, C. Villava, D.A. Griffith, A. Phillips, and J.C.W. Locke. 2018. Escherichia coli can survive stress by noisy growth modulation. Nat. Commun. 9: 5333.

30. Pedraza, J.M., and A. van Oudenaarden. 2005. Noise propagation in gene networks. Science. 307: 1965–1969.

31. Barr, A.R., S. Cooper, F.S. Heldt, F. Butera, H. Stoy, J. Mansfeld, B. Novák, and C. Bakal. 2017. DNA damage during S-phase mediates the proliferation-quiescence decision in the subsequent G1 via p21 expression. Nat. Commun. 8.

32. Reyes, J., J.-Y. Chen, J. Stewart-Ornstein, K.W. Karhohs, C.S. Mock, and G. Lahav. 2018. Fluctuations in p53 Signaling Allow Escape from Cell-Cycle Arrest. Mol. Cell. 71: 581–591.e5.

33. Shimoni, Y., S. Altuvia, H. Margalit, and O. Biham. 2009. Stochastic analysis of the SOS response in Escherichia coli. PLoS ONE. 4.

34. Verbruggen, P., T. Heinemann, E. Manders, G. von Bornstaedt, R. van Driel, and T. Höfer. 2014. Robustness of DNA Repair through Collective Rate Control. PLoS Comput Biol. 10: e1003438.

35. Nguyen, A.W., and P.S. Daugherty. 2005. Evolutionary optimization of fluorescent proteins for intracellular FRET. Nat. Biotechnol. 23: 355–360.

36. Wang, P., L. Robert, J. Pelletier, W.L. Dang, F. Taddei, A. Wright, and S. Jun. 2010. Robust growth of Escherichia coli. Curr. Biol. CB. 20: 1099–1103.

37. Landini, P., and M.R. Volkert. 1995. RNA polymerase alpha subunit binding site in positively controlled promoters: a new model for RNA polymerase-promoter interaction and transcriptional activation in the Escherichia coli ada and aidB genes. EMBO J. 14: 4329–4335.

38. Sedgwick, B. 1989. In vitro proteolytic cleavage of the Escherichia coli Ada protein by the ompT gene product. J. Bacteriol. 171: 2249–2251.

39. Saget, B.M., and G.C. Walker. 1994. The Ada protein acts as both a positive and a negative modulator of Escherichia coli’s response to methylating agents. Proc. Natl. Acad. Sci. 91: 9730–9734.

40. Li, G.-W., D. Burkhardt, C. Gross, and J.S. Weissman. 2014. Quantifying Absolute Protein Synthesis Rates Reveals Principles Underlying Allocation of Cellular Resources. Cell. 157: 624–635.

41. Bernstein, J.A., A.B. Khodursky, P.-H. Lin, S. Lin-Chao, and S.N. Cohen. 2002. Global analysis of mRNA decay and abundance in Escherichia coli at single-gene resolution using two-color fluorescent DNA microarrays. Proc. Natl. Acad. Sci. 99: 9697–9702.

42. Gillespie, D.T. 1977. Exact stochastic simulation of coupled chemical reactions. J. Phys. Chem. 81: 2340–2361.

43. Brewster, R.C., F.M. Weinert, H.G. Garcia, D. Song, M. Rydenfelt, and R. Phillips. 2014. The Transcription Factor Titration Effect Dictates Level of Gene Expression. Cell. 156: 1312–1323.

44. Yu, J., J. Xiao, X. Ren, K. Lao, and X.S. Xie. 2006. Probing gene expression in live cells, one protein molecule at a time. Science. 311: 1600–1603.

45. Paulsson, J. 2004. Summing up the noise in gene networks. Nature. 427: 415–418.

46. Peterson, J.R., J.A. Cole, J. Fei, T. Ha, and Z.A. Luthey-Schulten. 2015. Effects of DNA replication on mRNA noise. Proc. Natl. Acad. Sci. 112: 15886–15891.

47. Slager, J., and J.-W. Veening. 2016. Hard-Wired Control of Bacterial Processes by Chromosomal Gene Location. Trends Microbiol. 24: 788–800.

48. Alon, U. 2007. Network motifs: theory and experimental approaches. Nat. Rev. Genet. 8: 450–461.

49. Wilkinson, D.J. 2009. Stochastic modelling for quantitative description of heterogeneous biological systems. Nat. Rev. Genet. 10: 122–133.

50. Elowitz, M.B., A.J. Levine, E.D. Siggia, and P.S. Swain. 2002. Stochastic gene expression in a single cell. Science. 297: 1183–1186.

51. Cooper, S., and C.E. Helmstetter. 1968. Chromosome replication and the division cycle of Escherichia coli Br. J. Mol. Biol. 31: 519–540.

52. Dinant, C., M.S. Luijsterburg, T. Höfer, G. von Bornstaedt, W. Vermeulen, A.B. Houtsmuller, and R. van Driel. 2009. Assembly of multiprotein complexes that control genome function. J. Cell Biol. 185: 21–26.

53. Luijsterburg, M.S., G. von Bornstaedt, A.M. Gourdin, A.Z. Politi, M.J. Moné, D.O. Warmerdam, J. Goedhart, W. Vermeulen, R. van Driel, and T. Höfer. 2010. Stochastic and reversible assembly of a multiprotein DNA repair complex ensures accurate target site recognition and efficient repair. J. Cell Biol. 189: 445–463.

54. Taketomi, A., Y. Nakabeppu, K. Ihara, D.J. Hart, M. Furuichi, and M. Sekiguchi. 1996. Requirement for two conserved cysteine residues in the Ada protein of Escherichia coli for transactivation of the ada promoter. Mol. Gen. Genet. MGG. 250: 523–532.

